# Functional characterization of human Homeodomain-interacting protein kinases (HIPKs) in *Drosophila melanogaster* reveal both conserved functions and differential induction of HOX gene expression

**DOI:** 10.1101/2020.11.03.366393

**Authors:** Stephen D. Kinsey, Gerald A. Shipman, Esther M. Verheyen

## Abstract

Homeodomain-interacting protein kinases (Hipks) are a family of conserved proteins that are necessary for development in both invertebrate and vertebrate organisms. Vertebrates have four paralogues, Hipks 1-4. Mice lacking *Hipk1* or *Hipk2* are viable, however loss of both is lethal during early embryonic development, with embryos exhibiting homeotic skeletal transformations and incorrect HOX gene expression. While these results suggest Hipks have a role in regulating HOX genes, a regulatory mechanism has not been characterized, and further comparisons of the roles of Hipks in development has not progressed. One challenge with characterizing developmental regulators in vertebrates is the extensive redundancy of genes. For this reason, we used *Drosophila melanogaster*, which has reduced genetic redundancy, to study the functions of the four human HIPKs (hHIPKs). In *D. melanogaster*, zygotic loss of the single ortholog *dhipk* results in lethality with distinct eye and head defects. We used a *dhipk* mutant background to compare the ability of each hHIPK protein to rescue the phenotypes caused by the loss of dHipk. In these humanized flies, both hHIPK1 and hHIPK2 rescued lethality, while hHIPK3 and hHIPK4 only rescued minor *dhipk* mutant patterning phenotypes. This evidence for conserved functions of hHIPKs in *D. melanogaster* directed our efforts to identify and compare the developmental potential of hHIPKs by expressing them in well-defined tissue domains and monitoring changes in phenotypes. We observed unique patterns of homeotic transformations in flies expressing hHIPK1, hHIPK2, or hHIPK3 caused by ectopic induction of Hox proteins. These results were indicative of inhibited Polycomb-group complex (PcG) components, suggesting that hHIPKs play a role in regulating its activity. Furthermore, knockdown of PcG components phenocopied hHIPK and dHipk expression phenotypes. Together, this data shows that hHIPKs function in *D. melanogaster*, where they appear to have variable ability to inhibit PcG, which may reflect their roles in development.

**Author summary:** The redundancy of vertebrate genes often makes identifying their functions difficult, and *Hipks* are no exception. Individually, each of the four vertebrate *Hipks* are expendable for development, but together they are essential. The reason *Hipks* are necessary for development is unclear and comparing their developmental functions in a vertebrate model is difficult. However, the invertebrate fruit fly has a single essential *dhipk* gene that can be effectively removed and replaced with the individual vertebrate orthologs. We used this technique in the fruit fly to compare the developmental capacity of the four human *HIPKs* (*hHIPKs*). We found that *hHIPK1* and *hHIPK2* are each able to rescue the lethality caused by loss of *dhipk*, while *hHIPK3* and *hHIPK4* rescue minor patterning defects, but not lethality. We then leveraged the extensive adult phenotypes associated with genetic mutants in the fruit fly to detect altered developmental pathways when *hHIPKs* are mis-expressed. We found that expression of *hHIPKs 1-3* or *dhipk* each produce phenotypes that mimic loss of function of components of the Polycomb-group complex, which are needed to regulate expression of key developmental transcription factors. We therefore propose that *Hipks* inhibit Polycomb components in normal development, though details of this interaction remain uncharacterized.

## Introduction

Homeodomain-interacting protein kinases (HIPKs) are a family of conserved serine/threonine kinases that are necessary for development in both invertebrate and vertebrate organisms [1]. In *Drosophila melanogaster*, combined maternal and zygotic loss of the single homologue *hipk* (referred to hereafter as *dhipk*) results in early embryonic lethality, while zygotic loss alone results in larval and pupal lethality [2]. In vertebrates, which have four *Hipk* genes (*Hipks1-4*), experiments performed in mice show that knockout of individual *Hipk* genes is not lethal, however knockout of both *Hipk1* and *Hipk2* genes results in embryonic lethality, likely due to functional redundancy between the paralogues [3]. Interestingly, *Hipk1/2* double knockout mice share phenotypes with *D. melanogaster dhipk* knockout organisms, such as defects in eye and head structure and aspects of patterning and development [2–4].

While past research suggests Hipk1 and Hipk2 have similar developmental roles based on their apparent functional redundancy, a comparison with the highly similar Hipk3 or less similar Hipk4 proteins has not been comprehensively assessed. The kinase domain is the region of greatest similarity between human HIPK (hHIPK) paralogs, a similarity that extends to the orthologous dHipk (Fig 1A). Hipks share other structural features outside of the kinase domain that have been implicated in protein-protein interactions and regulating Hipk stability and localization, which have been reviewed by our group and others [1,5]. Individual knockouts of the four *Hipks* have been generated in mice by several groups, with each knockout producing distinct phenotypes that may be indicative of either divergent functions, different temporal-spatial expression, or both. For example, *Hipk1* knockout mice appear grossly normal, *Hipk2* knockout mice exhibit impaired adipose tissue development, smaller body size, and higher incidence of premature death, *Hipk3* knockout mice exhibit impaired glucose tolerance, and male *Hipk4* knockout mice are infertile due to abnormal spermiogenesis [6–10]. Unfortunately, these reported phenotypes come from a small number of sources focusing primarily on different tissues, so an exhaustive comparison of developmental roles for each Hipk is not possible using the current literature.

**Fig 1.**
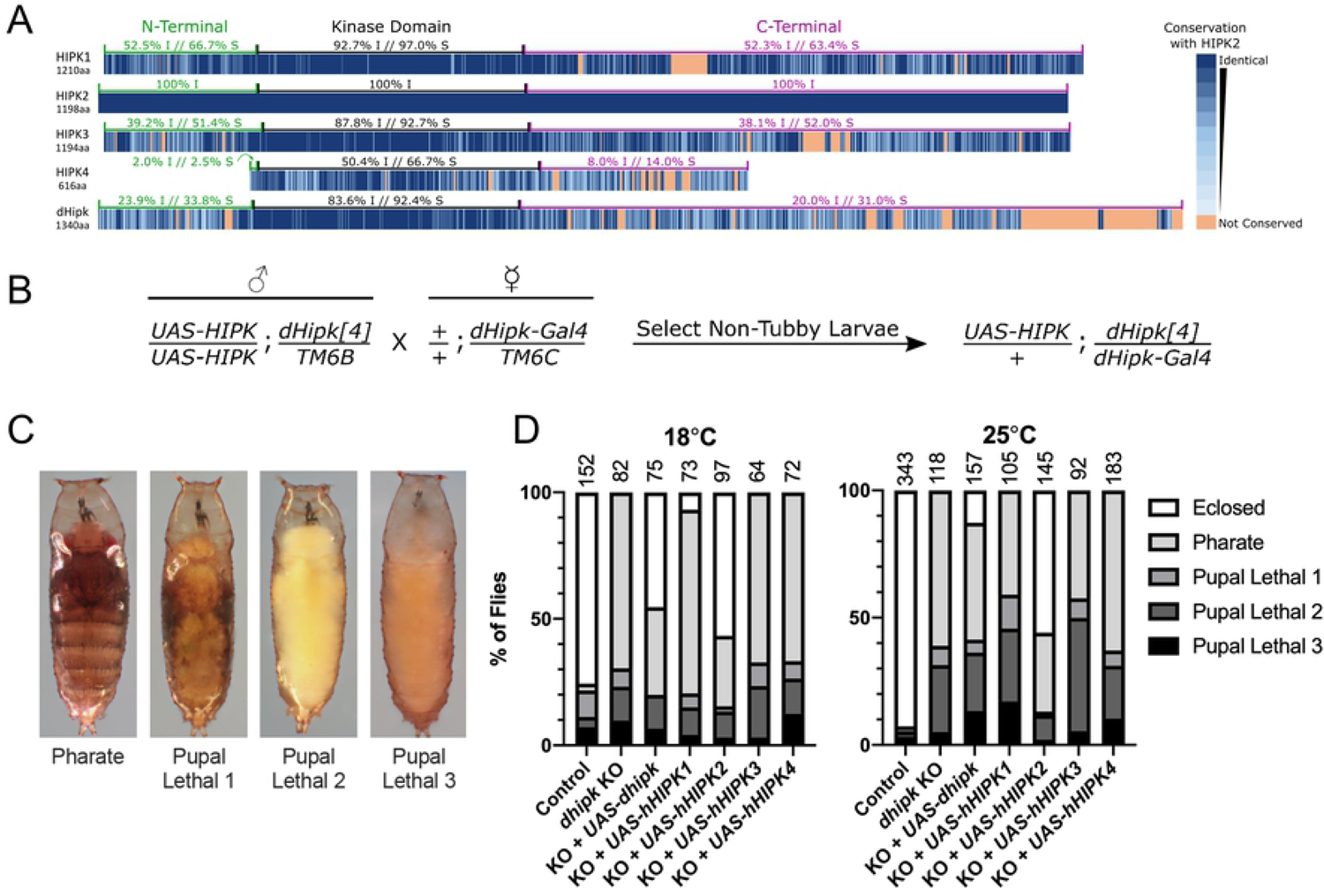
hHIPK1 and hHIPK2 rescue *dhipk* mutant lethality. (A) The four human HIPKs and the single dHipk protein amino acid sequences are each compared with hHIPK2, the most studied hHIPK, for amino acid identity and similarity. Dark blue indicates higher sequence similarity, while light blue indicates lower sequence similarity, and orange indicates lack of conservation between the protein and hHIPK2. Within each Hipk amino acid sequence, the kinase domain is the region of highest similarity when compared with hHIPK2. Less similarity is present in the N and C-terminal domains, where various interaction and regulatory domains exist, as reviewed by Rinaldo *et al*. (2008) and Schmitz *et al*. (2013). (B) The cross scheme used to generate *dhipk* mutant flies that expressed *UAS-hHIPKs* in the *dhipk* domain involved crossing two fly strains. A male fly homozygous for a *UAS-hHIPK* transgene on the 2^nd^ chromosome and heterozygous for the *dhipk[4]* mutant on the 3^rd^ chromosome over the balancer *TM6B* was crossed to a female fly with a wild-type 2^nd^ chromosome and homozygous for *dhipk-Gal4* on the 3^rd^ chromosome over the balancer *TM6C*. Resulting non-tubby progeny pupae were then scored for each cross. (C) The developmental stages were scored by assessing the pupal cases as described in the materials and methods. (D) Numbers at the top of the graph indicate the number of pupae scored per genotype. Flies were raised at the indicated temperatures with single-day egg lays. The furthest developmental stage of each pupae was recorded 5 days after control balancer flies eclosed and was plotted on the graph. Both male and female flies were combined for this experiment.

Vertebrate and *D. melanogaster* Hipks share conserved functions, as dHipk modulates signaling pathways important in normal development that are homologous to what various vertebrate Hipks interact with, including components of WNT, JNK, Hippo, and JAK/STAT signaling pathways [11–22]. Among Hipks, vertebrate Hipk2 in particular has been studied extensively for its role in responding to genotoxic stress, where it is stabilized upon lethal DNA damage and mediates p53-mediated cell death [23]. In fact, most studies involving vertebrate Hipk proteins focus on Hipk2, with few studies making comparisons with the highly similar Hipk1 and Hipk3. As of yet, no studies have assessed the functional equivalency of all vertebrate Hipks in development. Therefore, due to the similarity in known function between dHipk and vertebrate Hipks, and precedence for the study of human protein functions in *D. melanogaster* [24–26], we used the *D. melanogaster* model to compare the functions of the four hHIPKs. By expressing hHIPKs in both a *dhipk* knockout background, and in multiple tissues of a wild-type genetic background, we directly compared the developmental equivalence of the four hHIPKs under identical conditions. We uncovered previously unidentified functional similarities between hHIPKs in overall developmental potential, as well as unique differences when assessing their activity in developing epithelial tissues that form the adult wing, leg, and eye.

## Results

### hHIPK1 and hHIPK2 rescue *dhipk* mutant lethality

As a first step in characterizing hHIPKs in *D. melanogaster*, we tested whether expression of *hHIPKs* individually could rescue *dhipk* mutant phenotypes. To do this, we combined two existing *dhipk* mutant alleles to generate a transheterozygous (heteroallelic) knockout which gives rise to a severe zygotic loss of function phenotype (Lee *et al*., 2009). This knockout approach has three main benefits. First, it combines a *dhipk[4]* null allele which is missing the majority of *dhipk* exons, with a less-severe *dhipk-Gal4* knockin/knockout allele that disrupts endogenous *dhipk* expression while allowing expression of *UAS*-driven transgenes in the endogenous *dhipk* domain due to the insertion of Gal4 encoding sequences in the *dhipk* locus (S1 Fig). Second, this approach strongly decreases the amount *dhipk* mRNA (S2 Fig), without removing the maternal contribution. This is beneficial, since when maternal *dhipk* mRNA is removed, flies die at the embryonic stage, however mutants with a normal maternal *dhipk* mRNA contribution develop up to the late pupal stage, allowing for phenotypic analysis of fully developed adult tissues. Third, this approach reduces the effect of secondary mutations present in chromosomes carrying the individual *dhipk* mutant alleles that may contribute to lethality when made homozygous. In subsequent sections, *dhipk[4]*/*dhipk-Gal4* mutant flies are simply referred to as ‘*dhipk* mutants’.

As a proof of principle, we performed rescue experiments by expressing a UAS-controlled wildtype *dhipk* cDNA construct in the *dhipk* mutant background. We carried out rescue crosses at both 18° and 25°C to assay the effects of two levels of transgene expression, since the activity of Gal4 is enhanced at higher temperatures [27]. This was essential to determining optimal conditions, since our previous work has shown that overexpression of dHipk in a wildtype background causes numerous phenotypes including tumorigenic effects (Blaquiere and Wong *et al*., 2018; Wong, Liao and Verheyen, 2019; Wong *et al*., 2020). Crosses were set up at both temperatures and the degree of adult survival was determined (Fig 1B). In cases of pupal lethality, we quantified the stage at which lethality occurred based on morphology of the fly within their pupal case, as shown in Fig 1C. Control flies heterozygous for the *dHipk- Gal4/+* allele show ∼92% adult viability at 25°C and ∼75% adult viability at 18°C. Zero *dhipk* mutant flies eclosed at either temperature, with death occurring at various pupal stages as indicated in Fig 1D. We assessed the ability of transgenic *UAS-dhipk* to rescue this lethality. At 25°C when Gal4 has relatively high activity, *UAS-dhipk* rescued 12.7% of flies to eclosion, while at 18°C *UAS-dhipk* rescued 45.3% of flies to eclosion. We attribute the low rescue at the higher temperature to harmful effects caused by *dhipk* overexpression.

Having established our assay conditions, we tested the ability of the four *UAS-hHIPK* transgenes to rescue *dhipk* lethality (Fig 1D). Both *UAS-hHIPK1* and *UAS-hHIPK2* could rescue *dhipk* mutant flies to eclosion, though the degree to which they could do this varied. *UAS-HIPK1* rescued 6.8% of *dhipk* mutants at 18°C, while it was unable to rescue at 25°C. In contrast, HIPK2 rescued the lethality of ∼56% of *dhipk* mutants at both temperatures. Strikingly, this rescue was more effective than the rescue by *UAS-dhipk*. Neither *UAS-HIPK3* nor *UAS-HIPK4* rescued adult lethality.

### *hHIPKs* variably rescue *dhipk* mutant patterning phenotypes

While *UAS-hHIPK3* and *UAS-hHIPK4* do not rescue *dhipk* mutant lethality, it was unclear if these hHIPKs could rescue minor *dhipk* mutant patterning phenotypes observed in fully formed, yet inviable, pharate adults dissected from their pupal cases. *dhipk* mutant pharate adults have reduced compound eye size compared to wild-type and loss of the simple eyes, known as ocelli (Fig 2A-D) [2,31]. In addition, sensory bristles called macrochaetes that are anterior and posterior to the ocelli are lost in *dhipk* mutant pharate adults (Fig 2A, E, F). Combined, these are the most obvious external phenotypes of pharate *dhipk* mutant flies. Therefore, we assessed the ability of the *UAS-hHIPKs* to rescue the reduced eye size, and loss of ocelli and bristles. As with the *dhipk* mutant lethality rescue experiments, we carried out these crosses at both 18°C and 25°C (Figs 2, S3).

**Fig 2.**
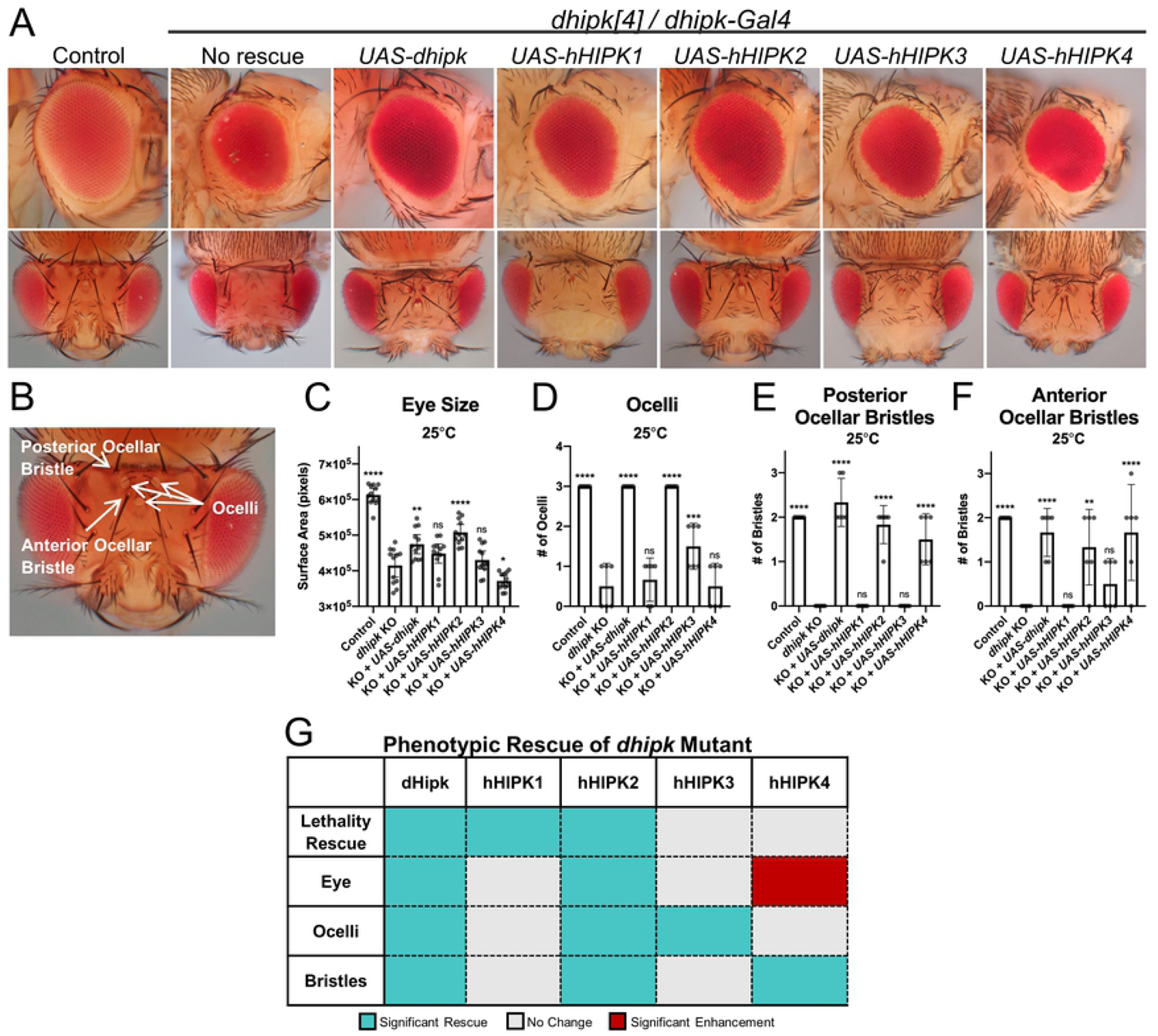
hHIPKs variably rescue minor *dhipk* mutant head phenotypes. (A) Representative heads and eyes from *dhipk* mutant flies expressing individual *UAS-hHIPKs* or *UAS-dHipk* using the *dhipk-Gal4* driver. (B) Location of the organs on the top of the head that were quantified in this figure. (C) The surface area of 12 eyes (6 flies) were imaged and measured for each cross. (D-F) The ocelli, posterior ocellar bristles, and anterior ocellar bristles of 6 heads were counted after imaging. (C-F) Comparisons in each graph are made to the *dhipk* mutant (*dhipk* KO) result. “Control” flies are of the genotype +/+ ; *dhipk-Gal4/+*. Error bars indicate the mean with a 95% confidence interval. A one-way ANOVA was performed followed by Dunnett’s test to correct for multiple comparisons for each dataset. P-values for the statistical analyses performed correspond to the following symbols: ≥0.0332 (ns), <0.0332 (*), <0.0021(**), <0.0002(***), < 0.0001(****). (G) Summary table of *dhipk* mutant rescue phenotypes. Only female flies were assessed for this experiment.

Overexpression of *UAS-dhipk* was able to significantly rescue each *dhipk* mutant phenotype when raised at 25°C and rescued all but the anterior bristle loss at 18°C (Figs 2A, C-F, S3). For the *UAS-hHIPKs*, only *UAS-HIPK2* could significantly rescue the reduced eye size (Fig 2B), while expression of *UAS-HIPK4* caused a significant decrease in eye size when compared to *dhipk* mutants at both temperatures (Figs 2C, S3A). The loss of ocelli in *dhipk* mutants was rescued by both *UAS-HIPK2* and *UAS-HIPK3*, but not by *UAS-HIPK1* or *UAS- HIPK4* when raised at either temperature (Fig 2D, S3B). None of the *UAS-hHIPKs* could significantly rescue the loss of posterior or anterior bristles in *dhipk* mutants at 18° (S3C, D Figs). In contrast, at 25° both *UAS-HIPK2* and *UAS-HIPK4* rescued the loss of both ocellar bristle pairs (Fig 2E, F). The ability of *UAS-dhipk* and *UAS-hHIPKs* to rescue *dhipk* mutant lethality, eye size, loss of ocelli, and loss of ocellar bristles are summarized in Fig 2G. Collectively these data revealed that within the same developmental context, the four hHIPKs can exert both shared and distinct effects that may reveal unique roles in development.

### hHIPK1 and 2 induce homeosis when expressed in wild-type *D. melanogaster* wings

The ability of *UAS-hHIPKs* to rescue impaired development in *dhipk* mutant flies suggests that hHIPKs expressed in *D. melanogaster* perform the same functions as dHipk. The varying ability of hHIPKs to rescue *dhipk* mutant flies may be due to divergent conserved functions, which could be observable in external *D. melanogaster* phenotypes. If so, the use of *D. melanogaster* tissues may provide us with a simple method of comparing developmental pathway alterations caused by the expression of hHIPKs. We therefore expressed hHIPKs in multiple wild-type tissues using the *dpp-Gal4* driver, which has well-defined expression patterns in the developing larval eye-antennal, wing, and leg imaginal discs [32]. To promote obvious phenotypic changes, these experiments were carried out at 29°C when Gal4 transcriptional activity is relatively high.

Expression of *UAS-hHIPK1* or *UAS-hHIPK2* causes notching of the adult wing when expressed using *dpp-Gal4, UAS-GFP* at 29°C (Fig 3A-C). The wing notching caused by *UAS- hHIPK1* expression is more pronounced than the phenotype caused by *UAS-hHIPK2*. Upon closer inspection, the region of the wing expressing either *UAS-hHIPK1* or *UAS-hHIPK2* contains small hairs and sensory bristles not normally found on the wing, instead resembling those found on halteres (Fig 3C). The altered development of a tissue causing it to fully or partially develop into another tissue is called homeotic transformation, or homeosis, and often occurs when key developmental regulators called homeotic (Hox) genes are dysregulated [33]. Halteres and wings are derived from similar larval tissues, the primary difference in their development being that haltere imaginal discs express the Hox gene *Ubx*, which inhibits Notch signaling at the dorsal-ventral boundary, while wing imaginal discs do not express Ubx [34]. Therefore, we stained 3^rd^ instar larval wing imaginal discs expressing the individual hHIPKs or dHipk to determine if ectopic Ubx expression was occurring. We found that expression of either hHIPK1 or hHIPK2 caused induction of Ubx in the wing pouch, but not in other wing imaginal disc regions where *dpp-Gal4* is expressed (Fig 3D). The degree of Ubx induction was greater in wing imaginal discs expressing hHIPK1 compared to those expressing hHIPK2, which matches the severity of the adult wing notching phenotypes. We also stained the same wing imaginal discs for Wingless (Wg) protein, which is a Notch target at the dorsal-ventral boundary of the wing pouch responsible for forming the edge of the wing. Since Notch signaling is inhibited by Ubx in the wing imaginal disc, we looked to see if Wg was decreased in response to *UAS-hHIPK* expression [34]. We found that wing imaginal discs expressing *UAS-hHIPK1* were missing Wg staining where *dpp-Gal4* intersects the dorsal-ventral boundary, while those expressing *UAS- hHIPK2* that induce lower levels of Ubx appeared to have intact Wg staining (S4 Fig). Together, this data suggests that hHIPK1 and hHIPK2 each induce Ubx expression in the wing pouch of wing imaginal discs, resulting in a wing-to-haltere homeotic transformation.

**Fig 3.**
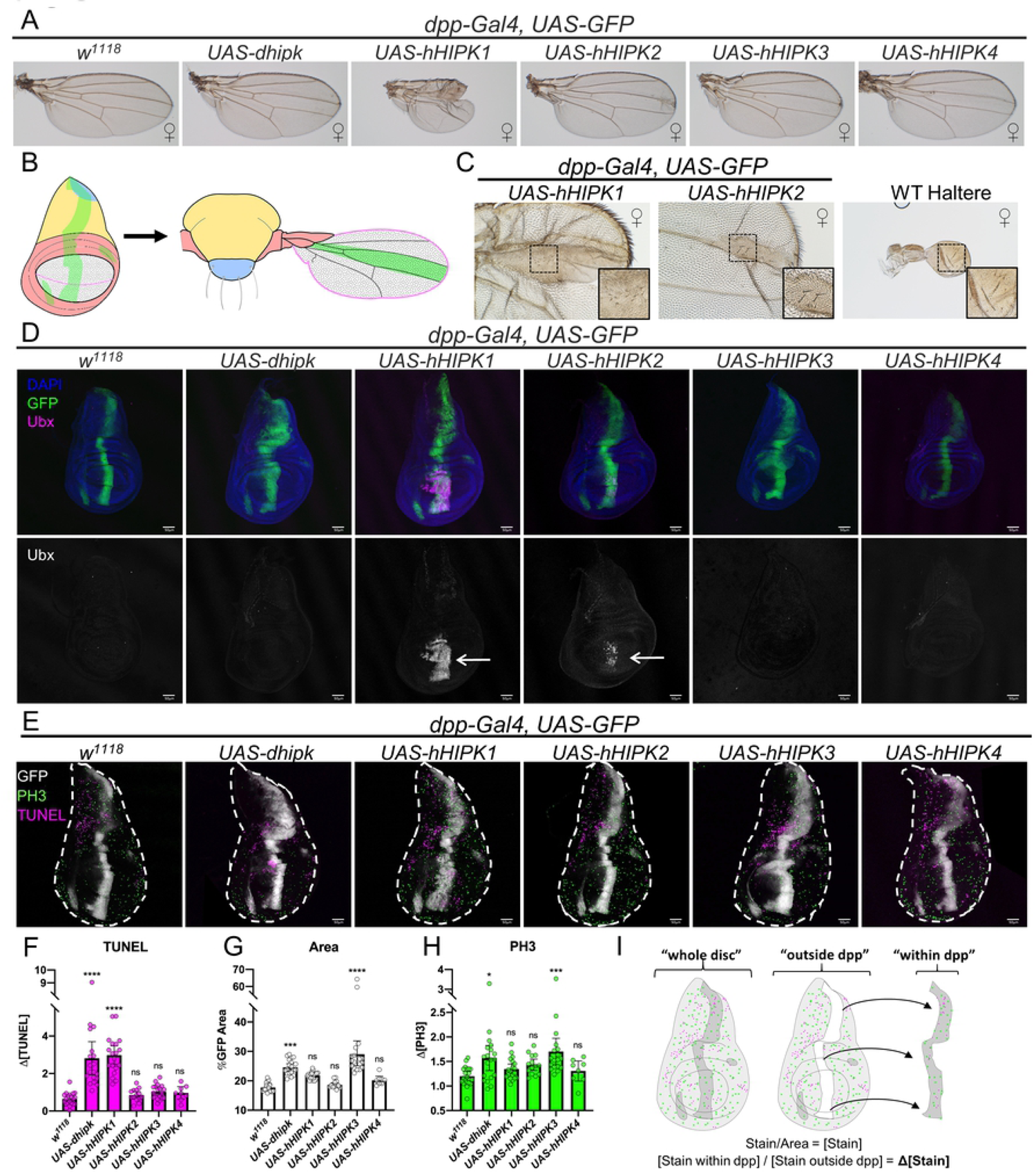
Effects of Hipk expression in wild-type *D. melanogaster* wings. (A) Representative adult wings dissected from the corresponding genotypes. (B) Graphical representation of the *dpp-Gal4* domain in larval wing disc and adult wing tissues. Green indicates the *dpp-Gal4* domain, while other colors and patterns indicate corresponding regions between the larval and adult wing. (C) Zoomed in image of *dpp-Gal4, UAS-hHIPK1 or UAS- hHIPK2* wing notching phenotype, compared to a wild-type haltere (images are to scale). Inset boxes for each image focus on similar phenotypes between the three images. (D) Representative images of late 3^rd^ instar imaginal wing discs dissected from larvae of the corresponding genotypes and stained for the Hox protein Ubx. Wing discs expressing *UAS-hHIPK1* or *UAS- hHIPK2* show Ubx induction in the wing pouch (arrows). Results were consistent across 10 wing imaginal discs assessed for each genotype. (E) Representative images of late 3^rd^ instar imaginal wing discs of the corresponding genotypes stained for mitotic marker PH3 and apoptosis marker TUNEL. (D, E) GFP marks the *dpp-Gal4* domain where *UAS* constructs are expressed. Scale bars are 50 µm. (F-H) Graphs plotting the change in TUNEL staining, Area, and PH3 staining, respectively, caused by expression of *UAS-Hipk* constructs. Error bars indicate the mean with a 95% confidence interval. A one-way ANOVA was performed followed by Dunnett’s test to correct for multiple comparisons for each dataset. P-values for the statistical analyses performed correspond to the following symbols: ≥0.0332 (ns), <0.0332 (*), <0.0021(**), <0.0002(***), < 0.0001(****). (I) Diagram explaining how changes in PH3 and TUNEL stains were quantified. For all images, the sex of the representative tissues was picked from mixed-sex samples unless otherwise noted by the female (♀) symbol. Crosses were performed at 29°C.

### hHIPKs variably induce cell death and proliferation

Wing notching can arise when cells making up the distal wing margin die [35–37]. Given that HIPKs have been implicated in promoting cell death under certain situations, we asked whether the notching is due to ectopic cell death [23,38–40]. We performed TUNEL staining in 3^rd^ instar wing imaginal discs to detect double stranded DNA breaks, which occur primarily in apoptotic cells [41]. Both *UAS-hHIPK1* and *UAS-hHIPK2* induce wing notching, and while *UAS-hHIPK1* expression did cause a significant increase in TUNEL staining, *UAS-hHIPK2* did not, suggesting that the cell death is not the primary cause of the wing notching phenotype (Fig 3E, F). Additionally, *UAS-dhipk* expression did cause a significant increase in TUNEL staining but did not produce the wing notching phenotype. Finally, while both *UAS-hHIPK1* and *UAS- dhipk* each caused a significant increase in TUNEL staining, the increased staining did not occur at the dorsal-ventral boundary of the wing pouch, which is the region that becomes the distal wing margin in the adult.

We have previously showed that using a different *UAS-dhipk* insertion strain (*UAS-Hipk*^*3M*^) that has higher expression levels than the *attP40* strain used in this work promotes cell proliferation in the wing imaginal disc [28–30]. Therefore, we tested the proliferative potential of each of the *UAS-hHIPKs* by measuring the size of the *dpp>GFP* expression domain after transgene expression, since increased proliferation would lead to more GFP-expressing cells. Expression of *UAS-dhipk or UAS-hHIPK3* each significantly increased the area of the *dpp* stripe in wing imaginal discs proportional to the size of the entire tissue (Fig 3E, G). To measure proliferation directly we stained wing imaginal discs for the mitotic marker phospho-histone 3 (PH3) and found that, similar to the results from measuring the *dpp-Gal4* expression area, expression of either *UAS-dhipk* or *UAS-hHIPK3* significantly increased cell proliferation in this tissue (Fig 3E, H). For both TUNEL and PH3 comparisons, the concentration of stain within the *dpp-Gal4* domain was measured both inside and outside of the main *dpp-Gal4* stripe. This data was used to calculate ratio of stain for each wing imaginal disc, which was then plotted to compare the genotypes (Fig 3I).We also expressed each of the *UAS-hHIPKs* or *UAS-dhipk* in eye-antennal imaginal discs using *ey-FLP*, which strongly drives *UAS* transgene expression in the entire tissue [42]. In this context, *UAS-hHIPK1, UAS-hHIPK3*, and *UAS-dhipk* each significantly increased the size of the eye-antennal imaginal discs, with the greatest increase found with hHIPK1 and hHIPK3, where obvious tissue distortions were also present (S5 Fig). Thus, we found that hHIPKs can variably induce proliferation in a tissue-dependent manner.

### hHIPK1 and 3 induce ectopic sex combs in male legs

Expressing *UAS-dhipk* at high levels using *dpp-Gal4* at 29°C causes malformed adult legs [30]. When expressing *UAS-hHIPKs* with *dpp-Gal4*, we found that *UAS-hHIPK3* caused similarly malformed legs, while *UAS-hHIPK1* caused less severe malformations (Fig 4A, S1 Table). Additionally, we found that *UAS-hHIPK1* and *UAS-hHIPK3* each caused ectopic sex comb formation on the middle and rear legs of males (Fig 4A, arrows, S1 Table). The *dpp-Gal4* domain is expressed in the region that produces sex combs in the leg-imaginal discs (Fig 4B). Because the ectopic sex comb phenotype is strongly associated with expression of the Hox protein Sex combs reduced (Scr, Fig 4C), we stained the larval imaginal discs that give rise to the middle legs with anti-Scr antibodies and found that those expressing *UAS-hHIPK1* or *UAS- hHIPK3* consistently showed ectopic Scr expression (Fig 4D).

**Fig 4.**
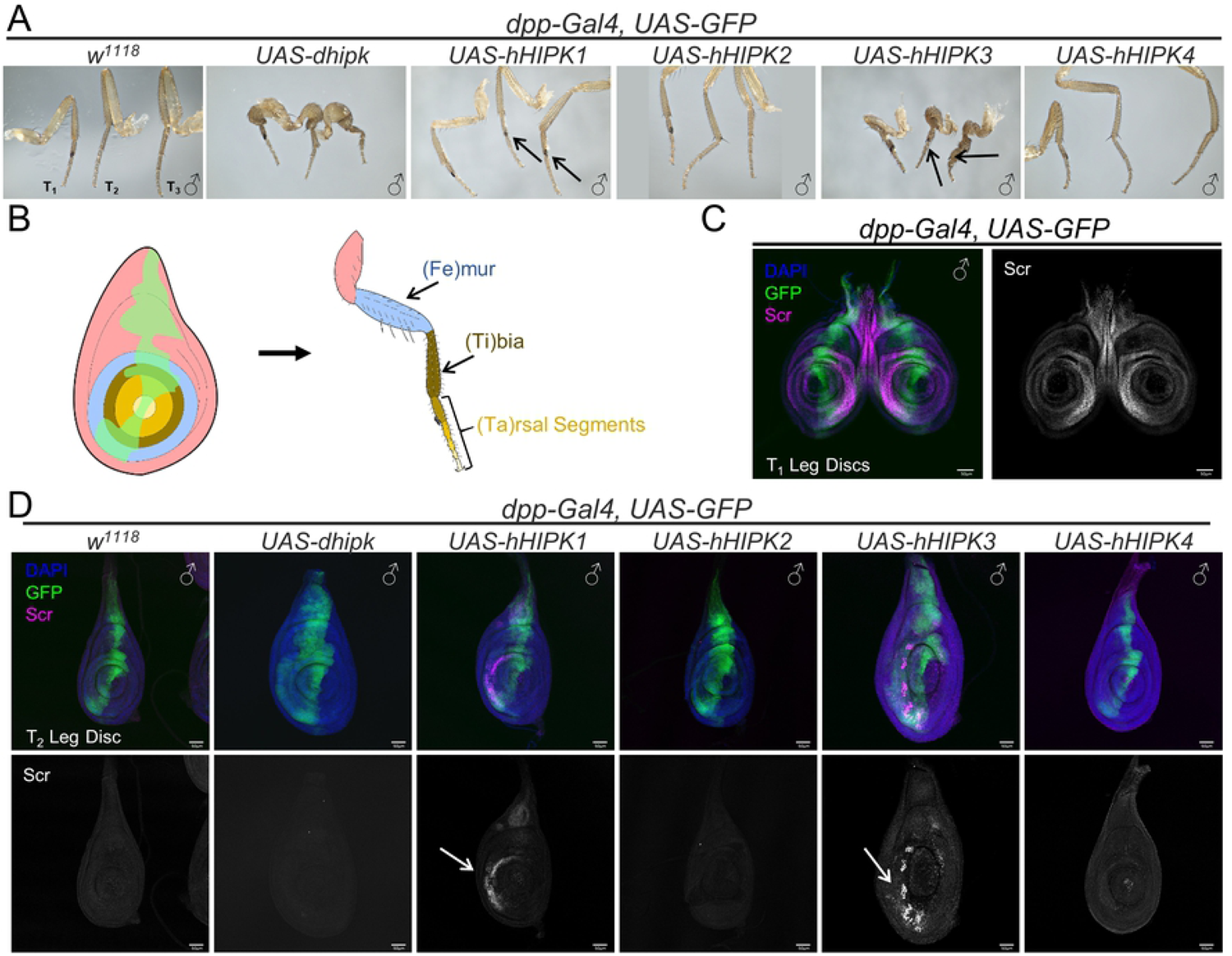
Effects of Hipk expression in wild-type *D. melanogaster* legs. (A) Adult legs dissected from the corresponding genotypes. Arrows indicate ectopic sex combs. (B) Graphical representation of the *dpp-Gal4* domain in larval leg imaginal disc and adult leg tissues. Green indicates the *dpp-Gal4* domain, while other colors indicate corresponding regions between the larval and adult leg. (C) Image of control late 3^rd^ instar T_1_ imaginal leg discs stained for the Hox protein Scr. (D) Representative images of late 3^rd^ instar T_2_ imaginal leg discs dissected from larvae of the corresponding genotypes and stained for the Hox protein Scr. Results were consistent across 10 T_2_ imaginal leg discs assessed for each genotype. GFP marks the *dpp-Gal4* domain where *UAS* constructs are expressed. All adult and larval flies assessed in this figure were male. Crosses were performed at 29°C. (C, D) Scale bars: 50µm.

### *UAS-dhipk* and *UAS-hHIPK1-3* expression phenocopies loss of Polycomb components

When staining for Ubx and Scr expression to determine a molecular cause for the adult wing and leg phenotypes, respectively, we also stained other larval tissues using each antibody. Ectopic expression of Ubx was detected in the wing pouch region of wing imaginal discs from larvae expressing *UAS-hHIPK1* or *UAS-hHIPK2*, but not in the leg or eye-antennal imaginal discs, and not from any imaginal discs expressing *UAS-hHIPK3, hHIPK4, or dhipk* (data not shown). Similarly, while ectopic Scr expression was detected in the middle and rear leg imaginal discs in flies expressing *UAS-hHIPK1* or *UAS-hHIPK3*, we did not observe Scr in the wing or eye-antennal disc, nor in any imaginal discs expressing *UAS-hHIPK2, hHIPK4*, or *dHipk* (data not shown). This tissue specific induction of Hox genes by hHIPKs is similar to what others have observed with Polycomb Group complex (PcG) mutants [43,44]. Mutations in *Polycomb* (*Pc*), a PcG component, have been shown to cause similar wing, leg, and antenna phenotypes as we observed with hHIPK1 expression [45–48]. *Pc* mutants are also known to mis-express Abdominal B (AbdB) in multiple tissues and developmental stages, including larval wing imaginal discs, adult ovaries, and embryos [49–51]. We therefore stained larval tissues expressing *UAS-hHIPKs* to detect AbdB and found that *UAS-hHIPK1* alone was able to induce ectopic AbdB expression in wing, leg, and eye-antennal imaginal discs (Fig 5A-C). Of note, the regions of tissue where AbdB was induced in wing or leg imaginal discs were different compared to the domains where Ubx or Scr, respectively, were induced by hHIPK1.

**Fig 5.**
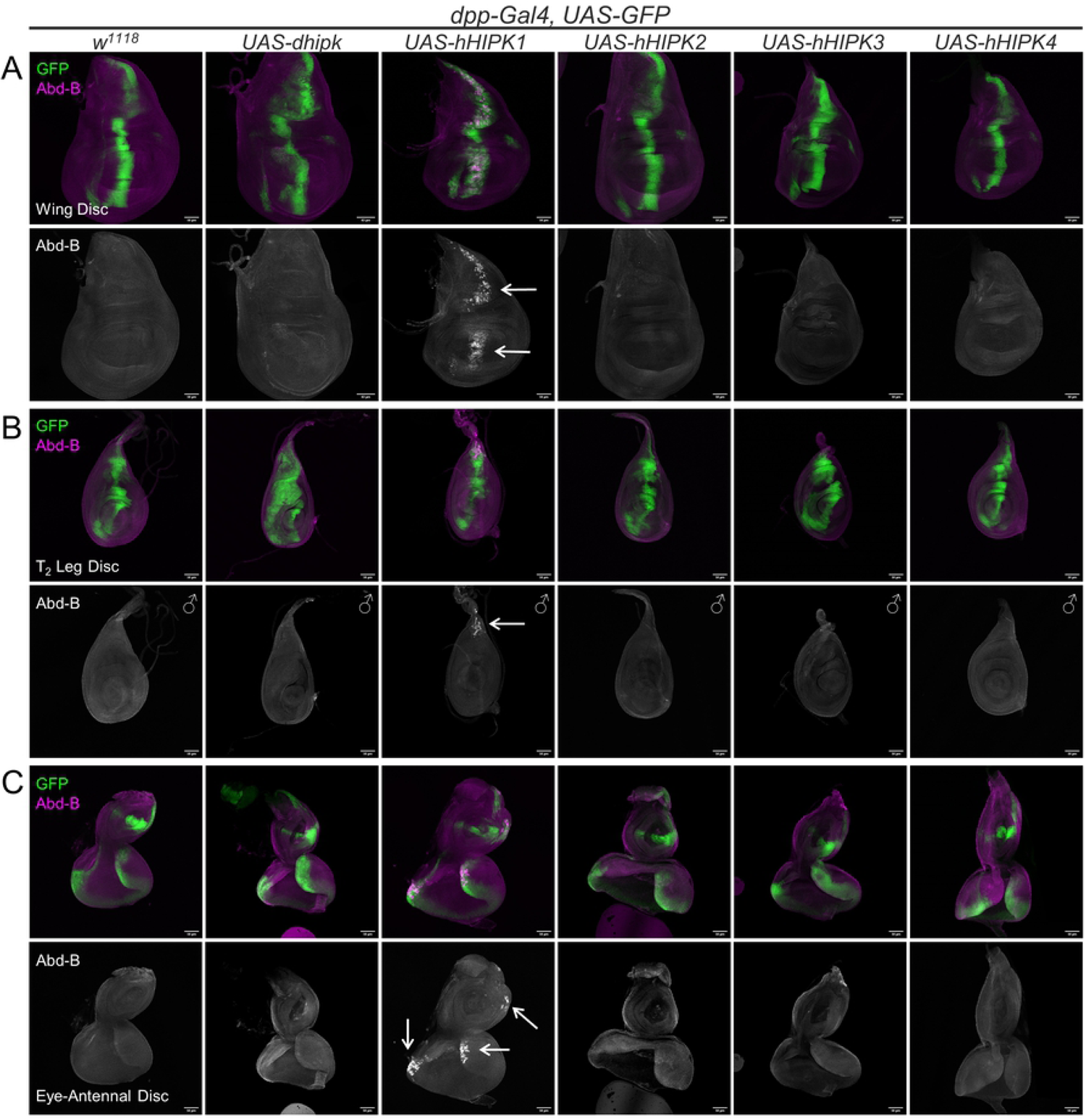
hHIPK1 induces Hox protein AbdB in wing, leg, and eye imaginal discs. Representative 3^rd^ instar imaginal (A) wing, (B) T_2_ leg, and (C) eye-antennal discs are shown for each of the corresponding crosses. Ectopic Abd-B staining caused by hHIPK1 is shown with arrows. Sex of the representative tissues are mixed unless otherwise noted by the male (♂) symbol. Crosses were performed at 29°C. Scale bars: 50µm.

**Fig 6.**
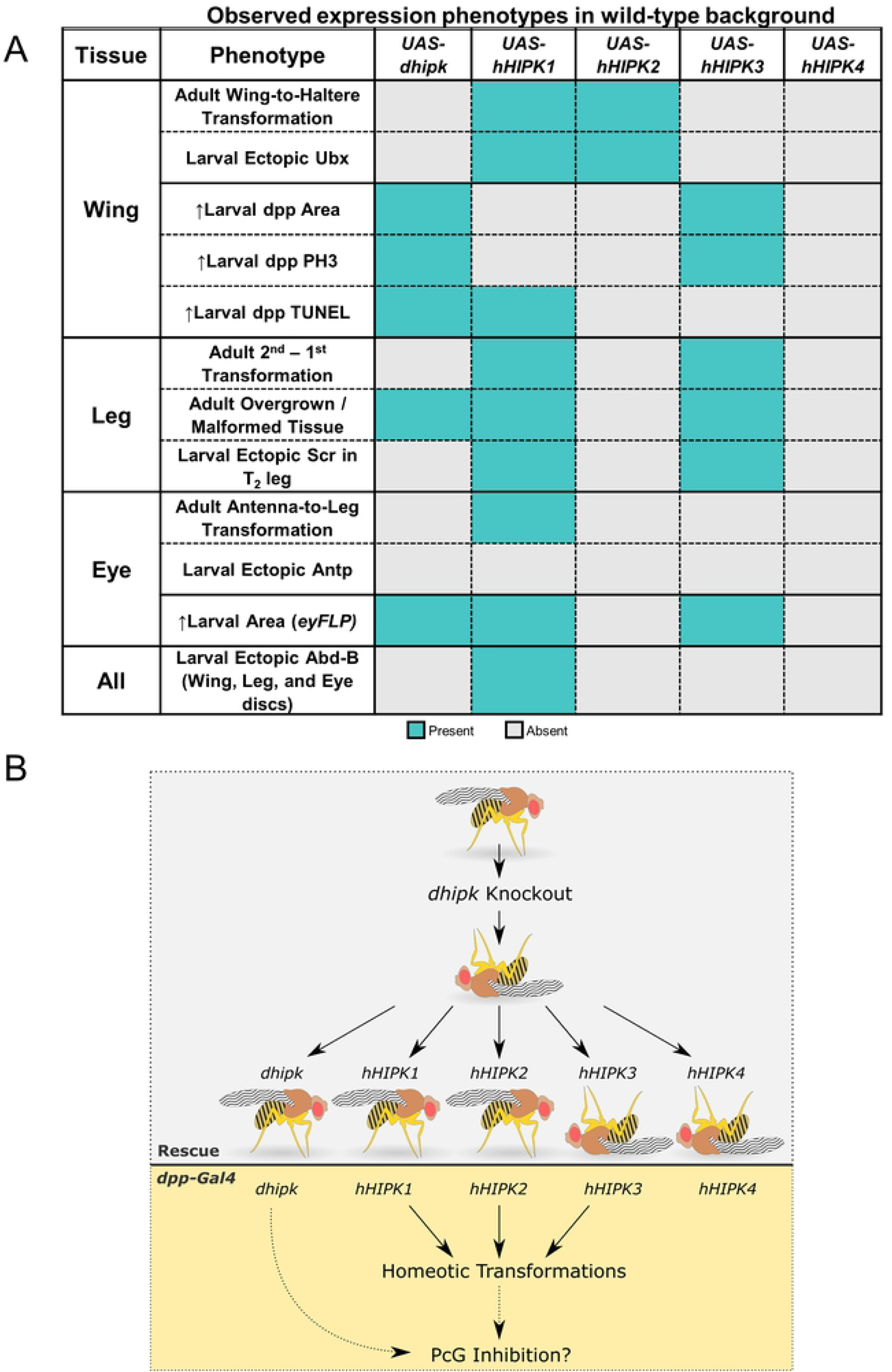
Summary of phenotypes induced by fly and human Hipks. (A) Summary table of phenotypes observed when Hipks are expressed at 29°C in non-mutant flies. Blue indicates a detected change, while grey indicates no change. All results in the table were achieved using *dpp-Gal4* except for the larval eye-antennal imaginal disc size (area) result, which utilized *eyFLP*. (B) Summary figure highlighting the main findings of this paper. Grey background indicates results from the rescue experiment, while yellow background indicates results from *dpp-Gal4* expression in a wild-type background

We associated the homeotic transformations observed with *UAS-hHIPK* expression with inactive PcG components, however *UAS-dhipk* did not produce an obvious homeotic transformation indicative of PcG alteration when over-expressed using *dpp-Gal4*. Given our finding that dHipk and hHIPKs have similar functions in the *dhipk* mutant rescue experiment, the lack of comparable homeotic transformation phenotypes in the *dpp-Gal4* experiment was surprising. Historically, the majority of phenotypes associated with altered PcG were obtained using genetic mutants for various PcG components, so it is possible that the overgrown/malformed leg phenotype that *UAS-dhipk* overexpression produced is due to reduced PcG activity but was not observed in PcG mutants due to impaired earlier development.

Therefore, to better assess the similarity of *UAS-dhipk* overexpression phenotypes to loss of PcG components, we used four *UAS-RNAi* lines targeting components of the two primary PcG complexes, PRC1 and PRC2, to reduce PcG activity in the *dpp-Gal4* domain. In addition to antenna-to-leg and wing-to-haltere transformations found with *Polycomb* (*Pc*), *Sex-combs-extra* (*Sce*), and *Enhancer of zeste* (*E(z)*) knockdown, we found that knockdown of, *Polyhomeotic* (*Ph- d*), *Sce*, or *E(z)* each produced overgrown/malformed adult legs (S6 Fig). While not an explicit homeotic transformation, these overgrown/malformed legs phenocopy the expression of *UAS- dhipk* (Figs 4A, S6).

## Discussion

Our results show that expressing human HIPKs 1-4 in *D. melanogaster* can substitute dHipk for many of its developmental functions. hHIPK1 and hHIPK2 are similar enough to dHipk that they each rescue lethality caused by mutant *dhipk*, while hHIPK3 and hHIPK4 are only capable of rescuing minor *dhipk* mutant patterning phenotypes. We also found that high-level expression of the hHIPKs in otherwise wild-type *D. melanogaster* tissues causes homeotic transformations indicative of PcG inhibition. Our work to compare the developmental functions and potential of the four human HIPKs under identical conditions builds upon work done by many groups to identify Hipk functions through knockout and overexpression studies in multiple organisms [1]. One of the primary motivations for this work was to make comparisons between HIPKs that were not possible in vertebrate models or cell culture experiments. Isono *et al*. (2006) demonstrated that Hipk1 and Hipk2 have overlapping, functionally redundant roles in mouse embryonic development, however it was not clear if other Hipks also shared this similarity [3]. Functional redundancy makes it difficult to study the functions of individual proteins in development, since the work necessary to make multiple strains of double or triple knockout mice is difficult, and sometimes impossible. It is much easier and financially practical to generate cell lines with multiple knockouts present, however in these you lose the perspective of the whole organism, which is key to identifying the necessity of these proteins in development. We thought the fly would be an excellent model to investigate this question of developmental necessity due to the strong similarity between dHipk and hHIPK protein sequences, and the simple and effective techniques we have available to knockout the single *dhipk* while expressing other hHIPKs in its place.

Our finding that hHIPK1 and hHIPK2 each rescue *dhipk* mutant lethality in flies suggests that the human HIPKs are functional in *D. melanogaster*, and that they share developmental roles both with each other, and with dHipk. This closely resembles mouse data from Isono *et al*., where Hipk1 and Hipk2 were shown to have overlapping roles during development by analysis of double Hipk1/Hipk2 knockouts [3]. There is no information about the possible functional redundancy between Hipk3 or Hipk4, so we assessed the extent of their functional similarities in our experimental model. The inability of hHIPK3 or hHIPK4 to rescue *dhipk* mutant lethality suggests their roles are more divergent from those of hHIPK1 and hHIPK2. This is not surprising for hHIPK4, since it lacks nearly all similarities to hHIPK1-3 and dHipk outside of the kinase domain, and even within the kinase domain it shares the least amount of similarity between them (see Fig 1). However, hHIPK3 shares nearly as much amino acid similarity with hHIPK2 as hHIPK1 does, so its inability to rescue *dhipk* mutant lethality may warrant further investigation into the significance of the amino acid sequence discrepancies between these proteins. The similarity between dHipk and hHIPK2 inferred from the strong *dhipk* mutant rescue by hHIPK2 also suggests that studies on dHipk may be used as a quick way to identify new hHIPK2 functions or targets.

While hHIPK1 rescues *dHipk* mutant lethality, it does not significantly rescue the patterning defects associated with *dhipk* mutants, unlike hHIPK2 which restores viability and patterning defects. In contrast, hHIPK3 and hHIPK4 rescue the loss of ocelli and loss of ocellar bristles in *dhipk* mutants, respectively, but not lethality. The limited depth of this analysis can only conclude that hHIPK3 and hHIPK4 may retain limited ancestral function or are so divergent that they rescue these phenotypes by some new mechanism. The comparison of protein sequence similarity, and previously published cellular localization data suggests that the normally nuclear hHIPK3 fits the former category, with cytoplasmic hHIPK4 fitting the latter. Similarly, the ability of hHIPK1 to rescue *dhipk* mutant lethality, but not patterning defects, may indicate functional divergence.

The ability of hHIPK1 and hHIPK2 to each rescue *dhipk* mutant lethality is strong evidence that hHIPKs were functioning correctly in *D. melanogaster*, however this experiment did not demonstrate what pathways were being modulated by hHIPK expression. Signaling pathway mutations are well characterized in *D. melanogaster*, to the extent that observation of distinct mutant phenotypes in adult cuticular structures often allows researchers to infer which signaling pathways or protein complexes are affected. Therefore, to reliably detect dHipk or hHIPK mediated changes in signaling, we needed to drive expression in well-defined regions of larval tissues that produce adult wing, leg, and head structures.

The well-established *dpp-Gal4* driver was selected for its common use among *D. melanogaster* researchers, and well-defined expression in multiple larval tissues, defined by co-expression of GFP. Expression of hHIPK1, hHIPK2, and hHIPK3 using *dpp-Gal4* each produced varied adult homeotic transformation phenotypes. hHIPK1 and hHIPK2 each caused wing-to- haltere transformations along with Ubx induction in the wing imaginal discs, while hHIPK3 and hHIPK4 had little to no effect on the adult wing structure and did not induce Ubx expression. Only wing imaginal discs expressing hHIPK1 showed a noticeable decrease in Wg staining at the dorsal-ventral boundary, suggesting that the lower level of Ubx induction caused by hHIPK2 was less able to decrease Notch signaling. hHIPK1 and hHIPK3 caused 2^nd^ and 3^rd^ legs to gain sex combs in males along with Scr induction in the corresponding leg imaginal discs, while hHIPK2 and hHIPK4 had no visible effect on these legs, nor did they induce ectopic Scr in leg imaginal discs. Finally, hHIPK1 alone was able to cause loss of aristae (S6 Fig), which is a minor antenna-to-leg transformation [52]. While the Hox protein Antp is frequently found to be ectopically expressed in eye-antennal imaginal discs that undergo antenna-to-leg transformations, we did not observe this (data not shown). However, partial antenna-to-leg transformations can occur without detectable levels of Antp [52].

Hipks are named for their initial discovery as binding partners of proteins containing homeodomains, so their ability to cause homeotic transformations may not seem to be a surprising result. While several studies have found direct protein-protein interactions between Hipks and homeodomain-containing proteins [53–55], it is important to note that the homeotic transformation phenotypes we have observed in response to hHIPK expression are not indicative of direct interaction with Hox proteins. Instead, the three homeotic transformations observed in these experiments are well characterized phenotypes associated with inactive PcG components, resulting in the upregulation of Hox gene transcription [56].

PcG components are broadly split into two protein complexes, Polycomb repressive complex 1 (PRC1) and PRC2, which act together to repress genes during development by regulating chromatin remodeling [56]. Previous research has shown that hHIPK2 can interact with and phosphorylate CBX4/Pc2, a PRC1 component homologous to *D. melanogaster* Polycomb (Pc), in the early response to DNA damage, however this interaction was found to promote transcriptional silencing in this context [57]. More recently, the same group has found that under otherwise normal conditions, targeting hHIPK2 to specific DNA regions causes de- condensation and de-repression of chromatin at those genomic loci [58]. The latter result supports our finding of normally repressed homeotic genes becoming de-repressed in response to hHIPK expression, however interactions between Hipks and PRC1 or PRC2 components that are important in the context of normal development have not yet been described. Given the strong homeotic transformations caused by hHIPK1, 2, and 3 presented in this research that are independent of DNA damage, we suspect that Hipks may guide development at least partially by inhibiting PcG components. An issue with this hypothesis is that dHipk overexpression did not produce any visible homeotic transformations like those produced by hHIPKs1-3. However, the leg deformities we observed in flies expressing dHipk, hHIPK1, and hHIPK3 are similar to what we observed when PcG components were knocked down with RNAi. Therefore, we suspect that dHipks are similarly inhibiting PcG, though more work needs to be done to detail these interactions, since the varying phenotypes caused by each hHIPK in *D. melanogaster* suggest that they are acting differently.

An alternative mechanism for how Hipks cause the various homeotic transformation phenotypes is by promoting the activity of proteins in the Trithorax group (TrxG) complex. TrxG proteins broadly act to promote gene expression by decreasing the compaction of chromatin, thereby increasing its accessibility [56]. Previous research has shown that expression of *D. melanogaster* TrxG component *trx* in the same *dpp-Gal4* domain used in this research produces nearly identical homeotic transformations as *UAS-hHIPK1* [52]. Because TrxG and PcG complex activities directly counter each other, it is difficult to assess whether these homeotic transformations are due to TrxG promotion or PcG inhibition. However, previous research describing Hipk/PcG interactions lead us to suspect that Hipks cause homeotic transformations through interactions with PcG components, not TrxG. Moving forward, we will investigate how the different Hipks alter chromatin, as well as clarifying the importance of this activity in development.

The pattern of Hox gene induction caused by HIPK expression is an important consideration. For example, the Hox protein Ubx is ectopically induced by hHIPK1 and hHIPK2, but only in the wing imaginal disc, not the leg or eye-antennal imaginal discs, and only in the wing pouch region, despite the domain of hHIPK expression being broader in this tissue. At the same time, the Hox protein AbdB is ectopically induced by hHIPK1 expression in both the wing pouch and the notum regions of the wing imaginal disc and is ectopically induced by hHIPK1 in other tissues. Clearly the Hox protein induction by hHIPKs is dependent on tissue region. The tissue-dependent response to hHIPK expression in these larval tissues highlights the importance of studying these proteins in a tissue context, rather than a cellular context, as the overall effect of HIPKs seems to vary depending on a cell’s existing developmental fate or pluripotency.

For the first time, all four vertebrate HIPKs have been assessed for their comparative developmental functions and potential under identical conditions. Together, these results show that hHIPK1 and hHIPK2 each function well enough in *D. melanogaster* to rescue lethality caused by mutant *dhipk*, the single fly Hipk homologue. Furthermore, hHIPK3 and hHIPK4 can rescue *dhipk* mutant patterning phenotypes. When expressed in domains that develop into adult cuticular structures, dHipk and hHIPKs1-3 each produce phenotypes that resemble loss of PcG components, suggesting that Hipks function to inhibit PcG components. This study collectively shows that Hipks share many conserved functions across species and validates the use of *D. melanogaster* as a tool to understand this complex and multi-facetted kinase family.

## Materials and methods

### Fly Stocks and Genetic Crosses

Previously described fly strains used in this work are **1:** *w*^*1118*^, **2:** *dhipk-Gal4* (*hipk[BG00855]*, BDSC #12779), **3:** *UAS-GFP* (BDSC #5431), **4:** *UAS-pc*^*RNAi*^ (BDSC #33964), **5:** *UAS-e(z)*^*RNAi*^ (BDSC #36068), **6:** *UAS-sce*^*RNAi*^ (BDSC #67924), **7:** *UAS-ph-d*^*RNAi*^ (BDSC #63018) **8:** *dhipk*^*4*^ [2], **9:** *dpp-Gal4/TM6B* [32], **10:** *UAS-HA-dhipk*^*attp40*^ [59], **11:** *eyFLP* ; *act>y*^*+*^*>Gal4, UAS-GFP* [42],. The details of how *UAS-myc-hHIPK1*^*attp40*^, *UAS-myc- hHIPK2*^*attp40*^, *UAS-myc-hHIPK3*^*attp40*^, and *UAS-myc-hHIPK4*^attp40^ were generated for this work is detailed in the section titled “Generation of plasmids and *UAS-hHIPK* fly stocks.” *dhipk* mutant rescue experiments were performed at 18°C and 25°C to determine the ideal Hipk expression levels by altering the abundance of Gal4-driven *UAS-Hipk* constructs, while experiments using *dpp-Gal4* were performed at 29°C to strongly increase *UAS-Hipk* expression. Flies were raised on standard media composed of 0.8g agar, 2.3g yeast, 5.7g cornmeal, and 5.2mL molasses per 100ml. “BDSC” is an acronym for the Bloomington Drosophila Stock Center.

### Terminology

As this study investigates human proteins expressed in *D. melanogaster*, it was necessary to clearly indicate which species of protein is specified in each experiment. Throughout this paper, *D. melanogaster* Hipk protein is written “dHipk” while mutants or DNA are referred to as *dhipk*, human HIPKs are written as “hHIPKs”, and in cases where reference is made to proteins from both species, “Hipks” is used.

### Generation of plasmids and transgenic *UAS-hHIPK* fly stocks

Plasmids containing the cDNA for human HIPKs were generously provided by two groups. Dr. Lienhard Schmitz gifted a plasmid containing *hHIPK1* isoform 1, and Dr. Seong- Tae Kim provided us plasmids containing *hHIPK3* isoform 2 and *hHIPK4*. The cDNA for *hHIPK2* isoform 1 was synthesized by GenScript® to match the NCBI reference sequence NM_022740.4. In cases where the gifted cDNAs did not exactly correspond to the translated NCBI reference protein sequences (NP_938009.1 for hHIPK1, NP_001041665.1 for hHIPK3, and NP_653286.2 for hHIPK4), we performed site-directed mutagenesis using the GeneArt™ Site-Directed Mutagenesis PLUS system to correct the cDNA sequence. The cDNAs that corresponded to these reference sequences were then tagged with N-terminal Myc-epitope tags before being cloned into a pUAST-attB backbone vector using NotI and XhoI restriction sites for *hHIPK1* and *hHIPK2*, BglII and KpnI sites for *hHIPK3*, and BglII and XhoI sites for *hHIPK4*. The four pUAST-attB-Myc-hHIPK plasmids were then sent to BestGene Inc. for injection into *D. melanogaster* embryos containing an attP40 site, allowing for stable integration to identical sites on the second chromosome. The resulting fly stocks each contain a single *Myc-hHIPK* cDNA under the control of a UAS promoter that is expressed in any cell expressing a Gal4 transcription factor.

### Adult *D. melanogaster* imaging and scoring rescue phenotypes

To quantify the stages of pupal lethality in the *dhipk* mutant rescue experiment, crosses were performed with 24-hour egg lays, and all non-Tubby pupal cases were collected 5 days after flies were expected to have eclosed. Pupal cases were scored into 5 categories: 1) “eclosed” flies were counted when pupal cases were empty. 2) Flies were scored as “pharate” when the adult head, thorax, and abdomen were fully developed and pigmented, but they were unable to eclose. 3) “Pupal lethal 1” was assigned to pupae that had defined head, thorax, and abdomen within the pupal case, but only had partial pigmentation. 4) “Pupal lethal 2” was assigned to pupae that had defined head, thorax, and abdomen, but no pigmentation. 5) “Pupal lethal 3” was assigned to pupae that had no defined head, thorax, or abdomen.

The pharate pupae and viable adults from the *dhipk* mutant viability rescue experiment were collected, and if necessary, gently removed from their pupal cases with dissecting tweezers before being immediately placed in 70% ethanol and stored at -20°C for preservation until they were photographed for the assessment and quantification of head phenotypes. Six randomly selected female flies from each cross were used for phenotype quantification. To image these flies, we used an 8-well BD Falcon CultureSlide (REF 354118) modified to have each well filled 1/3 with SYLGARD™ 184. Insect pins were bent at 90° and pinned into the solidified

SYLGARD so that the 90° bend was located near the top of the plastic well (S8 Fig). Immediately before imaging, flies were removed from 70% ethanol at -20°C to individual wells filled with 70% ethanol at room temperature and pinned to the planted insect pins while remaining submerged. The slides were then topped off with excess 70% ethanol before a coverslip was placed atop the wells. A resulting slide contained six female flies of the same genotype pinned at a stable position for imaging near the surface of the coverslip, while remaining submerged in ethanol. The ethanol was required to prevent flies drying out during imaging, and the coverslip was required to prevent vibrations on the surface of the ethanol that interfered with imaging. The same six flies were photographed three times to capture each eye (two images per fly) and the top of the head (one image per fly). Lighting was provided by an LED strip modified to encircle the CultureSlide, and a folded white tissue was placed under the CultureSlide to obtain a white/grey background.

Adult wings and legs were dissected in ethanol, then gently dried on a paper towel before being submerged in a small drop of Aquatex® (Sigma-Aldrich #1.08562) and covered in a coverslip. Small weights (EM stubs) were then placed on the coverslips while being heated to 60°C for 1 hour. All adult phenotypes were imaged using a Zeiss Axioplan2 microscope with an Optika C-P6 camera system.

### HIPK Protein Sequence Alignment

After confirming that our cDNA sequences correctly translated to the NCBI reference protein sequences for hHIPK1 isoform 1 (NP_938009.1), hHIPK2 isoform 1 (NP_073577.3), hHIPK3 isoform 2 (NP_001041665.1), hHIPK4 (NP_653286.2), and dHipk isoform A (NP_612038.2), each of the dHipk or hHIPK sequences were individually compared to hHIPK2 using the NCBI Multiple Sequence Alignment (MSA) tool [60]. On the MSA website, the Hipk that was being compared to hHIPK2 was set as the anchor. The FASTA alignment for this comparison was then downloaded and opened in Jalview (version 2.11.1.2) to extract the numerical conservation data between the Hipk in question and hHIPK2 [61]. The numerical conservation data (from 0 = no conservation, to 11 = identical amino acid) was then extracted and sent to Microsoft Excel (Excel 365), where numerical columns were converted to a color gradient. Each comparison to HIPK2 was lined up based on the location of the first conserved region. An image of the comparison was then exported as a PNG to Inkscape (version 0.92.4) for domain annotation, based on the NCBI annotation of the kinase domain.

### Immunocytochemistry and microscopy

Late third instar larval imaginal discs were dissected and stained using previously described methods [28]. The following primary antibodies were used: mouse anti-Ubx (1:50, DSHB Ubx FP3.38) mouse anti-Scr (1:50, DSHB anti-Scr 6H4.1), mouse anti-Abd-B (1:50, DSHB anti-ABD-B (1A2E9)), mouse anti-Antp (1:50, DSHB anti-Antp 4C3), mouse anti-Wg (1:50, DSHB 4D4), rabbit anti-PH3 Ser10 (1:500, Cell Signaling #9701S). Imaginal discs were imaged on a Zeiss LSM 880 using a dry 10x lens. Z-stacks were acquired, and images were processed in FIJI, where Z-stacks were converted to maximum intensity projections.

### PH3 and TUNEL assay quantification using wing imaginal discs

Dual PH3 and TUNEL assay staining was performed by first completing the normal wing disc dissection, fixing, washing, and primary antibody treatment protocol noted previously for PH3 (1:500 in block, Cell Signaling #9701S). Before secondary antibody staining, TUNEL staining was performed using the Roche *In Situ* Cell Death Detection Kit, TMR Red (Version 12, Cat. No. 12 156 792 910). Once the tissues were washed after the primary antibody treatment, the wash was removed, and 100ul of combined TUNEL assay components (92.7µL labelling solution + 8.3µL enzyme solution) was added to the tissues in a 1.6mL Eppendorf tube, along with 1:1000 goat α-rabbit fluorophore conjugated secondary antibody (Jackson ImmunoResearch, product # 711-605-152). The tissues were then incubated overnight (∼16 hours) on a rocker in the dark at 4°C. Staining regents were then removed, and samples were rinsed quickly with PBT before staining for 30 minutes with 1:500 DAPI solution. After DAPI staining, four more 10-minute washes were performed before wing discs were separated from other tissues and mounted in 70% glycerol on microscope slides. Wing imaginal discs were imaged as described in the previous section. Using FIJI [62–64], the area of the whole wing imaginal disc and *dpp-GFP* domains were measured, and PH3 or TUNEL positive cells were counted within each region automatically using the Analyze → Analyze Particles tool after thresholding. The change in concentration of PH3 or TUNEL positive cells between the dpp-GFP domain and the rest of the disc was then calculated.

### RNA extraction and qPCR

RNA extractions were performed using the Qiagen RNeasy® Plus Min Kit (#74134). RNA that was used to confirm reduced *dHipk* mRNA in *dhipk* mutant and rescue crosses, as well as verify the correct *hHIPK* expression in the rescue crosses, was collected from four combined wandering 3^rd^ instar larvae (two male and two female) for each cross. Larvae were washed in PBS before being spot dried on a clean paper towel and transferred to 300µl buffer RLT Plus, supplemented with freshly added β-mercaptoethanol to 1%. Larvae were homogenized with pestles by hand in 1.6mL tubes before being centrifuged for 3 minutes at maximum speed to pellet debris. Supernatant was transferred to a gDNA Eliminator spin column, with the remaining RNA extraction steps following the manufacturer’s instructions.

cDNA synthesis was performed using ABM® OneScript® Plus cDNA Synthesis Kit (#G236). For each sample, 100ng mRNA was used in combination with Oligo (dT) primers to perform first-strand cDNA synthesis of poly-adenylated mRNA following manufacturer’s instructions. Resulting cDNA was diluted 1:5 before being used for qPCR.

qPCR for each sample/primer mix was performed in triplicate with 10µl samples (technical replicates), utilizing Bioline’s sensiFAST SYBR Lo-ROX Kit (#BIO-94005) on an Applied Biosystems QuantStudio 3. 1µl of diluted cDNA was used per reaction. Primers targeting *rp49* were used as reference targets.

### Primers

*rp49* F: AGCATACAGGCCCAAGATCG

*rp49* R: TGTTGTCGATACCCTTGGGC

*dhipk* F: GCACCACAACTGCAACTACG

*dhipk* R: ACGTGATGATGGTGCGAACTC

*hHIPK1* F: GACCAGTGCAGCACAACCAC

*hHIPK1* R: GCCATGCTGGAAGGTGTAGG

*hHIPK2* F: GTCCACCAACCTGACCATGA

*hHIPK2* R: GGAGACTTCGGGATTGGCTA

*hHIPK3* F: GACATCAGCATTCCAGCAGC

*hHIPK3* R: GCTGTCTTCTGTGCCCAAAG

*hHIPK4* F: GCCTGAGAACATCATGCTGG

*hHIPK4* R: GCGACTGGATGTATGGCTCC

## Acknowledgements

We thank the following undergraduate students who participated in this research while training in the lab: Madeline Malczewska, Emerson Mohr, Justin Vinluan, and Rayna Brands. Stocks obtained from the Bloomington Drosophila Stock Center (NIH P40OD018537) were used in this study. The *eyFLP* stock was a gift from Amit Singh. We would like to acknowledge the Developmental Studies Hybridoma Bank for providing antibodies. We thank Drs. Lienhard Schmitz and Seong-Tae Kim for donating plasmids containing *hHIPK* cDNAs. We are grateful for the advice provided by Drs. Gritta Tettweiler and Don Sinclair on various aspects of this research. Also, we thank Z. Ding for help in generating the *UAS-HA-dHipk*^*attp40*^ plasmid.

---

**S1 Fig. Schematic of *dhipk* mutant allele generation**. (A) The *dhipk[4]* allele was generated by P-element excision, as described previously [2]. (B) The *dhipk-Gal4* allele was generated in the Baylor genetrap screen by insertion of a P-element containing a Gal4 exon into the beginning of the *dhipk* gene [65].

**S2 Fig. Validating *dhipk* knockout and *UAS-Hipk* expression using qPCR**. (A) The expression of *dhipk* was compared between wild-type, heterozygous *dhipk* mutant, transheterozygous *dhipk* knockout, and *dhipk* knockouts expressing *UAS-Hipks*. (B) Expression of specific *hHIPKs* or *dhipk* was confirmed in the respective *dhipk* mutant rescue experiments using qPCR. For each *UAS-hHIPK* or *UAS-dhipk* rescue assessed, *dhipk* transheterozygous knockouts were used as the control. (A,B) Two male and two female 3^rd^ instar larvae from each cross raised at 25°C were used in these experiments. Bars represent the mean, while error bars represent the upper and lower limits as defined by Quantstudio Design and Analysis Software.

**S3 Fig. Rescue of *dhipk* mutant patterning defects by hHIPKs at 18°C**. (A) The surface area of 12 eyes (6 flies) were imaged and measured for each cross. (B-D) The ocelli, posterior ocellar bristles, and anterior ocellar bristles of 6 heads were counted after imaging. (A-D) Comparisons in each graph are made to the *dhipk* mutant (*dhipk* KO) result. “Control” flies are of the genotype +/+ ; *dhipk-Gal4/+*. Error bars indicate the mean with a 95% confidence interval. A one-way ANOVA was performed followed by Dunnett’s test to correct for multiple comparisons for each dataset. P-values for the statistical analyses performed correspond to the following symbols: ≥0.0332 (ns), <0.0332 (*), <0.0021(**), <0.0002(***), < 0.0001(****).

**S4 Fig. Wg protein is reduced at the D/V boundary where *UAS-hHIPK1* is expressed**. 3^rd^ instar larval imaginal wing discs were stained as described in the materials and methods. Scale bars are 50µm. Flies were raised at 29°C.

**S5 Fig. Comparing the effects of *UAS-hHIPKs* and *UAS-dhipk* on eye-antennal disc size when expressed using *eyFLP***. The *eyFLP* genetic construct causes strong *UAS* transgene expression within the entire eye-antennal disc. (A) Representative images of eye-antennal imaginal discs from each cross, with *w*^*1118*^ used as the control. (B) Plotted data on the graph is from imaged eye-antennal discs with surface area measured using FIJI. Bars represent the mean, while error bars represent the 95% confidence interval. Scale bars are 50µm. Flies were raised at 29°C.

**S6 Fig. RNAi knockdown of PcG components phenocopies Hipk expression**. Fly stocks containing *UAS-RNAi* constructs expressed using *dpp-Gal4* causes homeotic transformation phenotypes similar to hHIPK1-3 expression (see Figs 3, 4, S7), and malformed legs similar to *dhipk* overexpression (see arrows). Flies were raised at 29°C..

**S7 Fig. Flies expressing *UAS-hHIPK1* in the eye-antennal disc do not develop aristae**. (A) Representative adult heads dissected from the corresponding genotypes. (B) Graphical representation of the *dpp-Gal4* domain in larval eye-antennal disc and adult head. Green indicates the *dpp-Gal4* domain, while other colors and patterns indicate corresponding regions between the larval and adult structures. Flies were raised at 29°C.

**S8 Fig. Container setup used to image adult flies in the *dhipk* mutant rescue experiments**. An 8-well Cultureslide was modified as described in the materials and methods to facilitate preparation of multiple samples for high-resolution imaging on a single slide.

**S1 Table.** Hipks variably induce leg deformities and ectopic sex combs. Representative flies are shown in Fig 4A. Front legs are listed as T_1_, middle legs as T_2_, rear legs as T_3_. The penetrance of leg deformities for each genotype is separated by leg section, where Fe indicates the Femur, Ti indicates the Tibia, and Ta indicates the Tarsal segments, as indicated in Fig 4B. Both leg distortion and sex comb frequencies are listed for males only. Female legs are distorted, but frequencies are not listed here. No female legs from these genotypes display ectopic sex combs. Flies were raised at 29°C.

| <u>Genotype</u> | <u># Legs Assessed</u> | <u>Deformed (by segment)</u> | <u>Sex Combs</u> |
| --- | --- | --- | --- |
| *dpp-Gal4, UAS-GFP*<br>*w<sup>1118</sup>* | T<sub>1</sub>: 7 / T<sub>2</sub>: 7 / T<sub>3</sub>: 7 | Fe: T<sub>1</sub>: 0% / T<sub>2</sub>: 0% / T<sub>3</sub>: 0%<br>Ti: T<sub>1</sub>: 0% / T<sub>2</sub>: 0% / T<sub>3</sub>: 0%<br>Ta: T<sub>1</sub>: 0% / T<sub>2</sub>: 0% / T<sub>3</sub>: 0% | T<sub>1</sub>: 100% / T<sub>2</sub>: 0% / T<sub>3</sub>: 0% |
| *dpp-Gal4, UAS-GFP*<br>*UAS-dhipk* | T<sub>1</sub>: 14 / T<sub>2</sub>: 14 / T<sub>3</sub>: 13 | Fe: T<sub>1</sub>: 100% / T<sub>2</sub>: 100% / T<sub>3</sub>: 100%<br>Ti: T<sub>1</sub>: 100% / T<sub>2</sub>: 100% / T<sub>3</sub>: 100%<br>Ta: T<sub>1</sub>: 0% / T<sub>2</sub>: 0% / T<sub>3</sub>: 0% | T<sub>1</sub>: 100% / T<sub>2</sub>: 0% / T<sub>3</sub>: 0% |
| *dpp-Gal4, UAS-GFP*<br>*UAS-hHIPK1* | T<sub>1</sub>: 21 / T<sub>2</sub>: 17 / T<sub>3</sub>: 19 | Fe: T<sub>1</sub>: 0% / T<sub>2</sub>: 0% / T<sub>3</sub>: 0%<br>Ti: T<sub>1</sub>: 0% / T<sub>2</sub>: 0% / T<sub>3</sub>: 0%<br>Ta: T<sub>1</sub>: 0% / T<sub>2</sub>: 0% / T<sub>3</sub>: 0% | T<sub>1</sub>: 100% / T<sub>2</sub>: 88.2% / T<sub>3</sub>: 78.9% |
| *dpp-Gal4, UAS-GFP*<br>*UAS-hHIPK2* | T<sub>1</sub>: 19 / T<sub>2</sub>: 19 / T<sub>3</sub>: 18 | Fe: T<sub>1</sub>: 80.0% / T<sub>2</sub>: 86.7% / T<sub>3</sub>: 94.4%<br>Ti: T<sub>1</sub>: 80.0% / T<sub>2</sub>: 73.3% / T<sub>3</sub>: 88.9%<br>Ta: T<sub>1</sub>: 20.0% / T<sub>2</sub>: 20.0% / T<sub>3</sub>: 38.9% | T<sub>1</sub>: 100% / T<sub>2</sub>: 0% / T<sub>3</sub>: 0% |
| *dpp-Gal4, UAS-GFP*<br>*UAS-hHIPK3* | T<sub>1</sub>: 15 / T<sub>2</sub>: 15 / T<sub>3</sub>: 18 | Fe: T<sub>1</sub>: 80.0% / T<sub>2</sub>: 86.7% / T<sub>3</sub>: 94.4%<br>Ti: T<sub>1</sub>: 80.0% / T<sub>2</sub>: 73.3% / T<sub>3</sub>: 88.9%<br>Ta: T<sub>1</sub>: 20.0% / T<sub>2</sub>: 20.0% / T<sub>3</sub>: 38.9% | T<sub>1</sub>: 100% / T<sub>2</sub>: 80.0% / T<sub>3</sub>: 44.4% |
| *dpp-Gal4, UAS-GFP*<br>*UAS-hHIPK4* | T<sub>1</sub>: 19 / T<sub>2</sub>: 19 / T<sub>3</sub>: 18 | Fe: T<sub>1</sub>: 0% / T<sub>2</sub>: 0% / T<sub>3</sub>: 0%<br>Ti: T<sub>1</sub>: 0% / T<sub>2</sub>: 0% / T<sub>3</sub>: 0%<br>Ta: T<sub>1</sub>: 0% / T<sub>2</sub>: 0% / T<sub>3</sub>: 0% | T<sub>1</sub>: 100% / T<sub>2</sub>: 0% / T<sub>3</sub>: 0% |

## References

1. Blaquiere JA, Verheyen EM. Homeodomain-Interacting Protein Kinases: Diverse and Complex Roles in Development and Disease. Current Topics in Developmental Biology. 2017. doi:10.1016/bs.ctdb.2016.10.002

2. Lee W, Andrews BC, Faust M, Walldorf U, Verheyen EM. Hipk is an essential protein that promotes Notch signal transduction in the Drosophila eye by inhibition of the global co-repressor Groucho. Dev Biol. Nov. 5, 20. 2009;325: 263–72. doi:10.1016/j.ydbio.2008.10.029

3. Isono K, Nemoto K, Li Y, Takada Y, Suzuki R, Katsuki M, et al. Overlapping roles for homeodomain-interacting protein kinases hipk1 and hipk2 in the mediation of cell growth in response to morphogenetic and genotoxic signals. Mol Cell Biol. 2006;26: 2758–71. doi:10.1128/MCB.26.7.2758-2771.2006

4. Inoue T, Kagawa T, Inoue-Mochita M, Isono K, Ohtsu N, Nobuhisa I, et al. Involvement of the Hipk family in regulation of eyeball size, lens formation and retinal morphogenesis. FEBS Lett. 2010;584: 3233–3238. doi:10.1016/j.febslet.2010.06.020

5. Rinaldo C, Siepi F, Prodosmo A, Soddu S. HIPKs: Jack of all trades in basic nuclear activities. Biochim Biophys Acta. 2008;1783: 2124–9. doi:10.1016/j.bbamcr.2008.06.006

6. Kondo S, Lu Y, Debbas M, Lin AW, Sarosi I, Itie A, et al. Characterization of cells and gene-targeted mice deficient for the p53-binding kinase homeodomain-interacting protein kinase 1 (HIPK1). Proc Natl Acad Sci U S A. 2003;100: 5431–6. doi:10.1073/pnas.0530308100

7. Sjölund J, Pelorosso FG, Quigley DA, DelRosario R, Balmain A. Identification of Hipk2 as an essential regulator of white fat development. Proc Natl Acad Sci U S A. 2014;111: 7373–8. doi:10.1073/pnas.1322275111

8. Chalazonitis A, Tang A a, Shang Y, Pham TD, Hsieh I, Setlik W, et al. Homeodomain interacting protein kinase 2 regulates postnatal development of enteric dopaminergic neurons and glia via BMP signaling. J Neurosci. 2011;31: 13746–57. doi:10.1523/JNEUROSCI.1078-11.2011

9. Shojima N, Hara K, Fujita H, Horikoshi M, Takahashi N, Takamoto I, et al. Depletion of homeodomain-interacting protein kinase 3 impairs insulin secretion and glucose tolerance in mice. Diabetologia. 2012;55: 3318–30. doi:10.1007/s00125-012-2711-1

10. Crapster JA, Rack PG, Hellmann ZJ, Le AD, Adams CM, Leib RD, et al. HIPK4 is essential for murine spermiogenesis. Elife. 2020;9. doi:10.7554/eLife.50209

11. Chen J, Verheyen EM. Homeodomain-interacting protein kinase regulates Yorkie activity to promote tissue growth. Curr Biol. 2012/07/31. 2012;22: 1582–6. doi:10.1016/j.cub.2012.06.074

12. Poon CLC, Zhang X, Lin JI, Manning SA, Harvey KF. Homeodomain-interacting protein kinase regulates Hippo pathway-dependent tissue growth. Curr Biol. 2012;22: 1587–94. doi:10.1016/j.cub.2012.06.075

13. Lan H-C, Li H-J, Lin G, Lai P-Y, Chung B. Cyclic AMP stimulates SF-1-dependent CYP11A1 expression through homeodomain-interacting protein kinase 3-mediated Jun N-terminal kinase and c-Jun phosphorylation. Mol Cell Biol. 2007;27: 2027–36. doi:10.1128/MCB.02253-06

14. Rochat-Steiner V, Becker K, Micheau O, Schneider P, Burns K, Tschopp J. FIST/HIPK3: a Fas/FADD-interacting serine/threonine kinase that induces FADD phosphorylation and inhibits fas-mediated Jun NH(2)-terminal kinase activation. J Exp Med. 2000;192: 1165–74. doi:10.1084/jem.192.8.1165

15. Hikasa H, Sokol SY. Phosphorylation of TCF proteins by homeodomain-interacting protein kinase 2. J Biol Chem. 2011/02/03. 2011;286: 12093–12100. doi:10.1074/jbc.M110.185280

16. Lee W, Swarup S, Chen J, Ishitani T, Verheyen EM. Homeodomain-interacting protein kinases (Hipks) promote Wnt/Wg signaling through stabilization of beta-catenin/Arm and stimulation of target gene expression. Development. 2008/12/18. 2009;136: 241–251. doi:10.1242/dev.025460

17. Louie SH, Yang XY, Conrad WH, Muster J, Angers S, Moon RT, et al. Modulation of the beta-catenin signaling pathway by the dishevelled-associated protein Hipk1. PLoS One. 2009;4: e4310. doi:10.1371/journal.pone.0004310

18. Shimizu N, Ishitani S, Sato A, Shibuya H, Ishitani T. Hipk2 and PP1c cooperate to maintain Dvl protein levels required for Wnt signal transduction. Cell Rep. 2014;8: 1391–404. doi:10.1016/j.celrep.2014.07.040

19. Swarup S, Verheyen EM. Drosophila homeodomain-interacting protein kinase inhibits the Skp1-Cul1-F-box E3 ligase complex to dually promote Wingless and Hedgehog signaling. Proc Natl Acad Sci U S A. 2011/06/02. 2011;108: 9887–92. doi:10.1073/pnas.1017548108

20. Hofmann TG, Stollberg N, Schmitz ML, Will H. HIPK2 regulates transforming growth factor-beta-induced c-Jun NH(2)-terminal kinase activation and apoptosis in human hepatoma cells. Cancer Res. 2003;63: 8271–7. Available: http://www.ncbi.nlm.nih.gov/pubmed/14678985

21. Hofmann TG, Jaffray E, Stollberg N, Hay RT, Will H. Regulation of homeodomain-interacting protein kinase 2 (HIPK2) effector function through dynamic small ubiquitin-related modifier-1 (SUMO-1) modification. J Biol Chem. 2005;280: 29224–29232. doi:10.1074/jbc.M503921200

22. Huang H, Du G, Chen H, Liang X, Li C, Zhu N, et al. Drosophila Smt3 negatively regulates JNK signaling through sequestering Hipk in the nucleus. Development. 2011/05/13. 2011;138: 2477–2485. doi:10.1242/dev.061770

23. D’Orazi G, Cecchinelli B, Bruno T, Manni I, Higashimoto Y, Saito S, et al. Homeodomain-interacting protein kinase-2 phosphorylates p53 at Ser 46 and mediates apoptosis. Nat Cell Biol. 2002;4: 11–19. doi:10.1038/ncb714

24. Link N, Bellen HJ. Using Drosophila to drive the diagnosis and understand the mechanisms of rare human diseases. Development. 2020;147. doi:10.1242/dev.191411

25. McGurk L, Berson A, Bonini NM. Drosophila as an In Vivo Model for Human Neurodegenerative Disease. Genetics. 2015;201: 377–402. doi:10.1534/genetics.115.179457

26. Ugur B, Chen K, Bellen HJ. Drosophila tools and assays for the study of human diseases. Dis Model Mech. 2016;9: 235–44. doi:10.1242/dmm.023762

27. Duffy JB. GAL4 system in Drosophila: a fly geneticist’s Swiss army knife. Genesis. 2002;34: 1–15. doi:10.1002/gene.10150

28. Blaquiere JA, Wong KKL, Kinsey SD, Wu J, Verheyen EM. Homeodomain-interacting protein kinase promotes tumorigenesis and metastatic cell behavior. Dis Model Mech. 2018;11. doi:10.1242/dmm.031146

29. Wong KKL, Liao JZ, Verheyen EM. A positive feedback loop between Myc and aerobic glycolysis sustains tumor growth in a Drosophila tumor model. Elife. 2019;8. doi:10.7554/eLife.46315

30. Wong KKL, Liu TW, Parker JM, Sinclair DAR, Chen YY, Khoo KH, et al. The nutrient sensor OGT regulates Hipk stability and tumorigenic-like activities in Drosophila. Proc Natl Acad Sci U S A. 2020. doi:10.1073/pnas.1912894117

31. Blaquiere JA, Lee W, Verheyen EM. Hipk promotes photoreceptor differentiation through the repression of Twin of eyeless and Eyeless expression. Dev Biol. 2014;390: 14–25. doi:10.1016/j.ydbio.2014.02.024

32. Staehling-Hampton K, Jackson PD, Clark MJ, Brand AH, Hoffmann FM. Specificity of bone morphogenetic protein-related factors: cell fate and gene expression changes in Drosophila embryos induced by decapentaplegic but not 60A. Cell Growth Differ. 1994;5: 585–93.

33. Pearson JC, Lemons D, McGinnis W. Modulating Hox gene functions during animal body patterning. Nat Rev Genet. 2005;6: 893–904. doi:10.1038/nrg1726

34. Weatherbee SD, Halder G, Kim J, Hudson A, Carroll S. Ultrabithorax regulates genes at several levels of the wing-patterning hierarchy to shape the development of the Drosophila haltere. Genes Dev. 1998;12: 1474–82. doi:10.1101/gad.12.10.1474

35. Brun S, Rincheval-Arnold A, Colin J, Risler Y, Mignotte B, Guénal I. The myb-related gene stonewall induces both hyperplasia and cell death in Drosophila: rescue of fly lethality by coexpression of apoptosis inducers. Cell Death Differ. 2006;13: 1752–62. doi:10.1038/sj.cdd.4401861

36. Adachi-Yamada T, Fujimura-Kamada K, Nishida Y, Matsumoto K. Distortion of proximodistal information causes JNK-dependent apoptosis in Drosophila wing. Nature. 1999;400: 166–169. doi:10.1038/22112

37. Giraldez AJ, Cohen SM. Wingless and Notch signaling provide cell survival cues and control cell proliferation during wing development. Development. 2003;130: 6533–43. doi:10.1242/dev.00904

38. Link N, Chen P, Lu WJ, Pogue K, Chuong A, Mata M, et al. A collective form of cell death requires homeodomain interacting protein kinase. J Cell Biol. 2007;178: 567–574. doi:10.1083/jcb.200702125

39. Hofmann TG, Möller A, Sirma H, Zentgraf H, Taya Y, Dröge W, et al. Regulation of p53 activity by its interaction with homeodomain-interacting protein kinase-2. Nat Cell Biol. 2002;4: 1–10. doi:10.1038/ncb715

40. Li X, Zhang R, Luo D, Park S-J, Wang Q, Kim Y, et al. Tumor necrosis factor alpha-induced desumoylation and cytoplasmic translocation of homeodomain-interacting protein kinase 1 are critical for apoptosis signal-regulating kinase 1-JNK/p38 activation. J Biol Chem. 2005;280: 15061–70. doi:10.1074/jbc.M414262200

41. Gavrieli Y, Sherman Y, Ben-Sasson SA. Identification of programmed cell death in situ via specific labeling of nuclear DNA fragmentation. J Cell Biol. 1992;119: 493–501. doi:10.1083/jcb.119.3.493

42. Pagliarini RA, Xu T. A genetic screen in Drosophila for metastatic behavior. Science. 2003;302: 1227–31. doi:10.1126/science.1088474

43. Busturia A, Morata G. Ectopic expression of homeotic genes caused by the elimination of the Polycomb gene in Drosophila imaginal epidermis. Development. 1988;104: 713–20.

44. Beuchle D, Struhl G, Müller J. Polycomb group proteins and heritable silencing of Drosophila Hox genes. Development. 2001;128: 993–1004.

45. Lewis EB. A gene complex controlling segmentation in Drosophila. Nature. 1978;276: 565–70. doi:10.1038/276565a0

46. Denell RE. Homoeosis in drosophila. II. A genetic analysis of polycomb. Genetics. 1978;90: 277–289.

47. Hannah-Alava A. Developmental Genetics of the Posterior Legs in Drosophila Melanogaster. Genetics. 1958;43: 878–905.

48. Lindsley DL, Grell EH. Genetic variations of Drosophila melanogaster. Carnegie Institute of Washington Publication. 1968.

49. Simon J, Chiang A, Bender W. Ten different Polycomb group genes are required for spatial control of the abdA and AbdB homeotic products. Development. 1992;114: 493–505.

50. Zhu J, Ordway AJ, Weber L, Buddika K, Kumar JP. Polycomb group (PcG) proteins and Pax6 cooperate to inhibit in vivo reprogramming of the developing Drosophila eye. Development. 2018;145. doi:10.1242/dev.160754

51. Gandille P, Narbonne-Reveau K, Boissonneau E, Randsholt N, Busson D, Pret A-M. Mutations in the polycomb group gene polyhomeotic lead to epithelial instability in both the ovary and wing imaginal disc in Drosophila. PLoS One. 2010;5: e13946. doi:10.1371/journal.pone.0013946

52. Sadasivam DA, Huang D-H. Maintenance of Tissue Pluripotency by Epigenetic Factors Acting at Multiple Levels. Schwartz YB, editor. PLOS Genet. 2016;12. doi:10.1371/journal.pgen.1005897

53. Kim YH, Choi CY, Lee SJ, Conti MA, Kim Y. Homeodomain-interacting protein kinases, a novel family of co-repressors for homeodomain transcription factors. J Biol Chem. 1998;273: 25875–9. doi:10.1074/jbc.273.40.25875

54. Kim EA, Noh YT, Ryu M-J, Kim H-T, Lee S-E, Kim C-H, et al. Phosphorylation and transactivation of Pax6 by homeodomain-interacting protein kinase 2. J Biol Chem. 2006;281: 7489–97. doi:10.1074/jbc.M507227200

55. Steinmetz EL, Dewald DN, Walldorf U. Homeodomain-interacting protein kinase phosphorylates the Drosophila Paired box protein 6 (Pax6) homologues Twin of eyeless and Eyeless. Insect Mol Biol. 2018;27: 198–211. doi:10.1111/imb.12363

56. Kassis JA, Kennison JA, Tamkun JW. Polycomb and Trithorax Group Genes in Drosophila. Genetics. 2017;206: 1699–1725. doi:10.1534/genetics.115.185116

57. Roscic A, Möller A, Calzado MA, Renner F, Wimmer VC, Gresko E, et al. Phosphorylation-Dependent Control of Pc2 SUMO E3 Ligase Activity by Its Substrate Protein HIPK2. Mol Cell. 2006. doi:10.1016/j.molcel.2006.08.004

58. Haas J, Bloesel D, Bacher S, Kracht M, Schmitz ML. Chromatin Targeting of HIPK2 Leads to Acetylation-Dependent Chromatin Decondensation. Front Cell Dev Biol. 2020;8. doi:10.3389/fcell.2020.00852

59. Tettweiler G, Blaquiere JA, Wray NB, Verheyen EM. Hipk is required for JAK/STAT activity during development and tumorigenesis. PLoS One. 2019;14. doi:10.1371/journal.pone.0226856

60. Multiple Sequence Alignment Viewer [Internet]. Bethesda (MD): National Library of Medicine (US), National Center for Biotechnology Information; [cited 16 May 2020]. Available: https://www.ncbi.nlm.nih.gov/projects/msaviewer/

61. Waterhouse AM, Procter JB, Martin DMA, Clamp M, Barton GJ. Jalview Version 2--a multiple sequence alignment editor and analysis workbench. Bioinformatics. 2009;25: 1189–91. doi:10.1093/bioinformatics/btp033

62. Schindelin J, Arganda-Carreras I, Frise E, Kaynig V, Longair M, Pietzsch T, et al. Fiji: an open-source platform for biological-image analysis. Nat Methods. 2012;9: 676–82. doi:10.1038/nmeth.2019

63. Schindelin J, Rueden CT, Hiner MC, Eliceiri KW. The ImageJ ecosystem: An open platform for biomedical image analysis. Mol Reprod Dev. 2015;82: 518–29. doi:10.1002/mrd.22489

64. Schneider CA, Rasband WS, Eliceiri KW. NIH Image to ImageJ: 25 years of image analysis. Nat Methods. 2012;9: 671–5. doi:10.1038/nmeth.2089

65. Bellen HJ, Levis RW, He Y, Carlson JW, Evans-Holm M, Bae E, et al. The Drosophila gene disruption project: Progress using transposons with distinctive site specificities. Genetics. 2011. doi:10.1534/genetics.111.126995

